# Small RNA-guided transgene repression systems enable toxic gene cloning in bacteria

**DOI:** 10.64898/2026.08.18.745554

**Authors:** Anna Pratt, Jeffrey M. Staub

## Abstract

Multiple vectors and bacterial strains have been developed to enable cloning and amplification of DNA plasmids used in bioengineering applications when transgenic components are toxic to the host. These include plasmids that limit readthrough transcription into transgenic sequences and host strains carrying mutations to minimize recombination or plasmid copy number. However, these techniques are insufficient in cases where transgene expression elements are recognized by the bacterial transcriptional apparatus, or the translation products have functions in cellular metabolism. Here we demonstrate two platforms that mitigate bacterial expression of transgenes driven by the prokaryotic-like promoters of chloroplast transgenes destined for use in plant plastid genetic engineering applications. Both an engineered CRISPRi approach and utilization of the native *E. coli* Hfq repression system resulted in significant knockdown of plasmid-borne transgene expression, resulting in reproducibly successful cloning and plasmid amplification. The advancements reported here will facilitate synthetic biology studies generally, and enable complex transgenic studies in prokaryotic-like organelles.

## INTRODUCTION

The production of plasmid-based vectors destined for transformation of plant or animal cells is often limited by the inability to clone the component parts into the *Escherichia coli* K12 (*E. coli*) production host (Sleight et al., 2010). Cloning failures most often arise from metabolic overload during plasmid propagation due to unintended transcription and translation of recombinant genes or regulatory elements that are active in the bacterial host. These unintended effects can disrupt essential cellular processes, interfere with membrane integrity, protein folding, or redox balance. In response, host cells selectively favor plasmid variants that reduce this metabolic burden, driving the accumulation of mutations that alleviate toxic transgenic expression. Cloning artifacts include DNA sequence polymorphisms in transgene protein-coding regions or their expression signals, insertion of transposable elements into transgene expression cassettes, unwanted recombination or deletion of construct features, or cloning assembly failure.

The host’s metabolic burden is particularly pronounced during high-copy plasmid replication and scale-up, where the cumulative expression of transgene products amplifies stress responses. Repression of transgene expression during plasmid production would allow for improvement of cloning outcomes. Several plasmid- and host-based transgene repression systems have been developed and are commercially available (reviewed for example in de Lorenzo and Martinez-Garcia, 2025; Kaur et al., 2018; Saida et al., 2006; Silva et al., 2012). These systems are designed to prevent unwanted transcriptional or translational readthrough from plasmid vector sequences into recombinant expression cassettes or to control the copy number of the recombinant plasmid to limit transgenic protein accumulation. However, a significant gap in the effectiveness of these cloning systems exists when the transgene expression signals are recognized and active in bacteria or recombinant proteins themselves have biochemical functions in the bacterial host.

Plant and animal cells harbor organelles (mitochondria and plastids) that evolved from ancient eubacterial endosymbionts. These organelles have retained many features of the ancient endosymbiont, most notably, their bacterial-like gene expression signals (Alberts et al., 2002; Liere et al., 2011; Small et al., 2013; Tadini et al., 2020). For example, mitochondrial genes in plants and animals have retained promoter signals recognized by a nuclear-encoded single subunit-type RNA Polymerase (NEP) with similarity to bacteriophage T7 RNA Polymerase, that has been transferred from the organelle genome to the host during evolution. In addition, plant plastids utilize a second, multi-subunit plastid-encoded polymerase (PEP) that recognizes consensus bacterial sigma 70-like promoter sequences. Due to their significant similarity to bacterial expression systems, there is currently no means of repressing the expression of mitochondrial and plastid transgenes during their cloning in bacterial hosts. As a result, transgene cassette mutations accumulate during cloning attempts, causing significant delays or the need to abort transformation vector production, significantly impacting research and/or commercial timelines of transgenic animals and plants.

Here, we describe the development of two modular transgene repression systems targeting expression elements recognized in *E. coli* that will result in toxic overexpression. Each system is based on a recently discovered bacterial small-RNA guided repression mechanism (Xie et al., 2020; Zhang et al., 2021). We show first that the bacterial-derived CRISPR interference (CRISPRi) system that relies on guide RNA targeting of a “dead” Cas nuclease to halt transcription by binding to its target DNA sequence and preventing initiation or translocation of RNA Polymerase (Price et al., 2020). Secondly, we developed a method of gene repression based on the Hfq protein (Na et al., 2013; Yoo et al., 2013) and its partner small synthetic RNA (sRNA) to silence the expression of target transgenes. The sRNA is designed to bind the transgenic mRNA and recruit binding of endogenous *E. coli* Hfq protein, preventing translation of targeted sequences. While these gene silencing systems have recently been used to study specific endogenous bacterial genes (Chen et al., 2015; Yang et al., 2018; Zhang et al., 2020), no universal cloning vectors or bacterial hosts utilizing these systems are yet commercially available.

As proof-of-concept toward development of universal cloning systems, we target repression of plastid transgene expression signals that drive extraordinarily high level of recombinant protein expression in *E*.*coli*. The results described here show reduction of recombinant protein expression 15 - 65%. Both the CRISPRi- and Hfq-based repression systems were efficacious as a single copy integrated into the *E. coli* genome while sequences targeted for repression are carried on multicopy plasmids. Furthermore, we show a near elimination of sequence polymorphisms and structural rearrangements during cloning of vectors capable of toxic levels of recombinant protein expression compared to current commercially available *E. coli* strains. Bacterial strains with these transgene repression components integrated into the genome separate the repression apparatus from recombinant plasmids that are amplified and purified from the host, simplifying the deployment and use of these approaches for standard cloning practices. Thus, the transgene repression systems reported here are a new tool that broadly enables cloning of toxic genes in bacteria.

## RESULTS

### CRISPR interference (CRISPRi) approach to plasmid-borne toxic transgene repression

#### Rhamnose-inducible dMad7

The first transgene repression system tested is based on dead Mad7 (dMad7), a Cas12a-type nuclease, that was created specifically for CRISPRi (Price et al., 2020). Like other Cas nucleases, Mad7 can exhibit toxicity in *E. coli* when over-expressed (Mund et al. 2023). Therefore, we placed expression of dMad7 under a rhamnose inducible promoter, P_rha_ (Giacalone et al., 2006), to allow fine-tuning of expression and reduce any potential toxic effects in *E. coli*.

Expression strength and rhamnose inducibility of the P_rha_ promoter was first tested using *E. coli* DH10B cells harboring plasmid pP_rha_404 (P_rha_404; Fig. 1) that carries a P_rha_::GFP transgene cassette on a high-copy number plasmid. As can be seen in Fig. 1, GFP fluorescence is near background levels in P_rha_404 cells in the absence of rhamnose. In contrast, addition of 1mM rhamnose to an overnight bacterial culture increased GFP fluorescence to ∼0.5 X 10^5^ RFU, confirming the strong activity and inducibility of the P_rha_ promoter. Interestingly, a control strain carrying plasmid pPTS389 that expresses GFP from the soybean plastid 16S rRNA gene promoter (GmPrrn) and bacteriophage T7 gene 10 (G10L; Olins et al., 1988) translational control sequence (GmPrrn_G10L) accumulates about twice as much GFP fluorescence as the rhamnose-induced P_rha_404 strain, illustrating the exceptionally strong promoter activity of the plastid gene promoter and G10L in *E. coli*.

**Figure 1.**
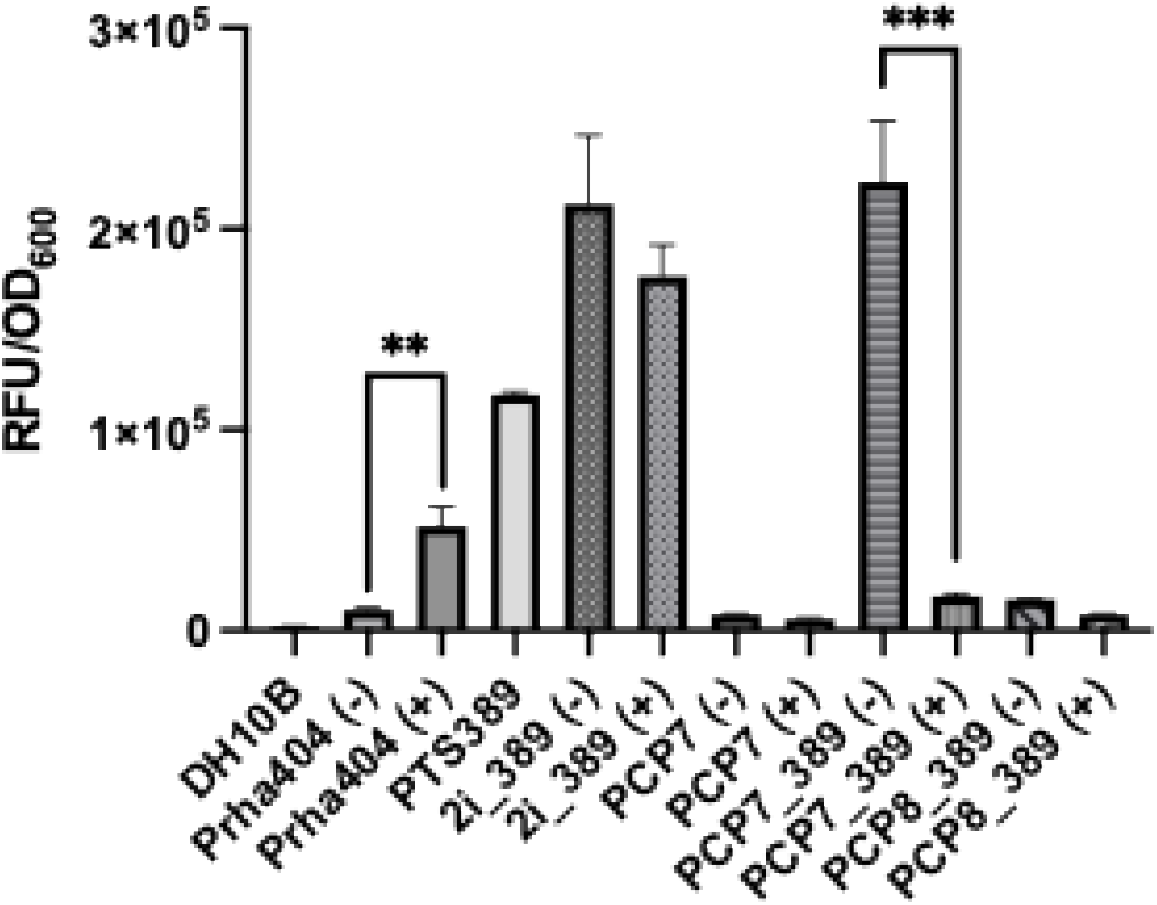
GFP fluorescence levels in bacterial cells carrying plastid transgenes and CRISPRi repression elements. Strains were grown overnight and then assayed for GFP activity, reported as relative fluorescence units (RFU) normalized to culture density (OD_600_). DH10B and the PTS389 strain were grown without rhamnose and are included as negative and positive controls, respectively. The PCP2i_389 (2i389) control strain has an integrated dMad7 but no gRNAs. PCP7 contains integrated dMad7 and gRNA but does not contain GFP. (-) indicates absence of rhamnose, (+) indicates the addition of rhamnose. Stars indicate statistical differences between cultures (N=3).

#### Integration of dMad7 and guide RNAs into the *E. coli* genome

Separation of the transgene repression apparatus from plasmid-borne sequences enables modular cloning, amplification and purification of plasmids for downstream applications. Therefore, dMad7 and guide RNAs (gRNAs) were integrated into the *E. coli* genome at the phage φ80 and λ *attB* sites, respectively, using the CRIM system of Haldimann and Wanner (2001) as described in Methods.

The P_rha_::dMad7 expression cassette was integrated into the *E. coli* φ80 *attB* site, as shown in Fig. 2A. Integration of the intact P_rha_::dMad7 expression cassette was confirmed by sequencing of the locus. Although an *E. coli* insertion element was found between the antibiotic resistance marker used for strain selection and the P_rha_::dMad7 expression cassette, the insertion element did not disrupt the dMad7 expression cassette. This strain was then transformed with a λ integrase helper plasmid to facilitate subsequent integration of gRNA sequences, creating strain PCP6 that was used for downstream experiments (Fig. 2B).

**Figure 2.**
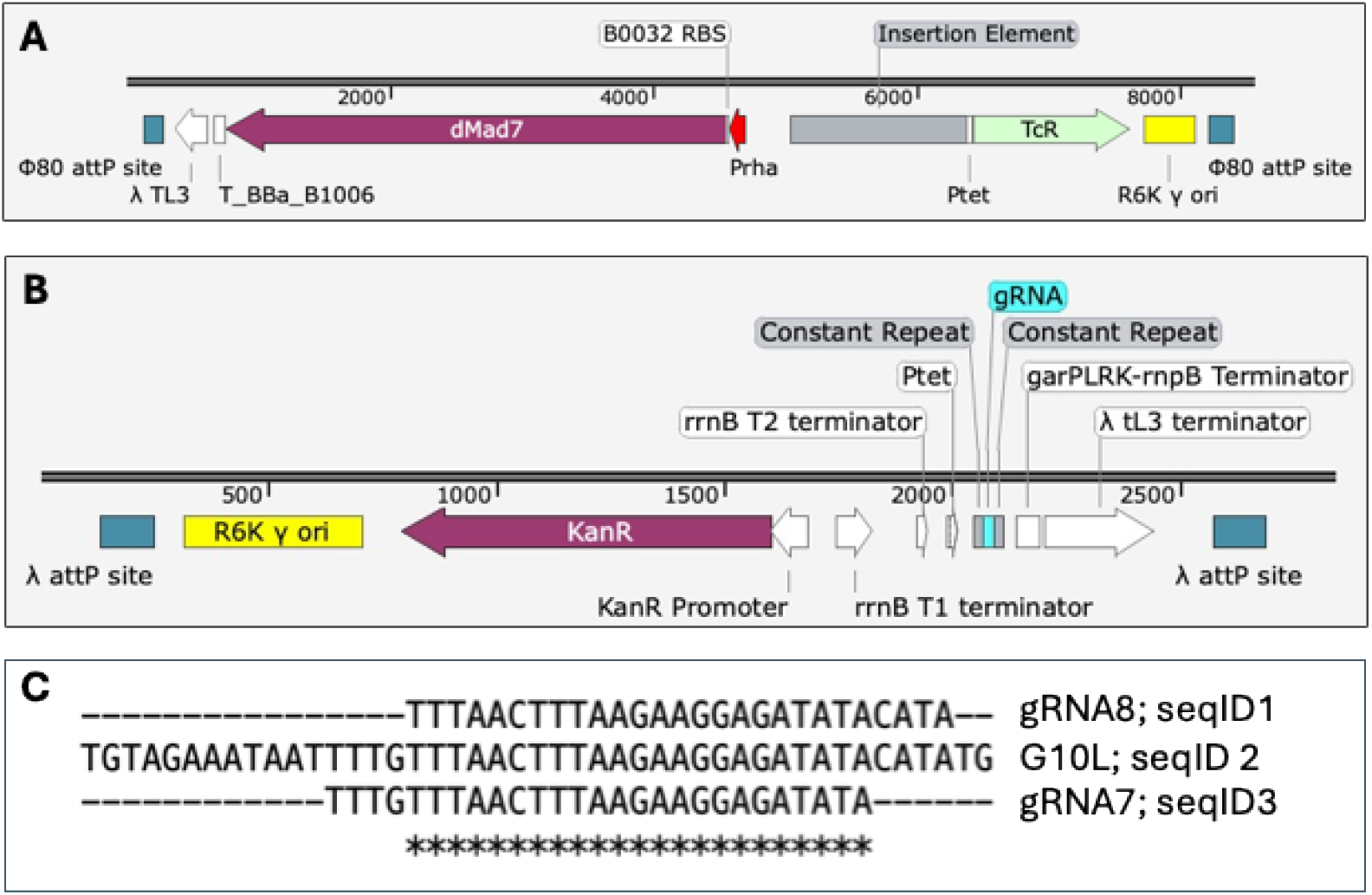
*E. coli* genomic loci showing dMad7 and guide RNA integration sites, and guide RNA sequences complimentary to the bacteriophage T7 G10 leader (G10L). A) Integration into the phage φ80 site of the rhamnose-inducible dMad7 cassette was selected via a tetracycline resistance (TcR) selectable marker. Note the position of the cointegrated *E. coli* insertion element. R6K ori and λ TL3 derived from the pPCP2 transforming plasmid; Ptet promoter drives TcR gene; B0032 ribosome binding site and T_BBa_B1006 terminator from iGEM. B) Integration into the λ attP site of the gRNA sequences flanked by the constant repeat sequence was selected via kanamycin resistance (KanR) selectable marker. Ptet promoter driving the gRNA; garPLRK-rnpB, rrnB T1 and T2 terminators. C). Alignment of the gRNA sequences to the target G10L. Asterisks indicate nucleotide identities.

#### gRNA targets

The G10L enhances translation of plastid transgenes by up to 100-fold (Ye et al. 2001) and is often used to overexpress recombinant proteins in chloroplasts (for example, Gray et al., 2009; Staub et al. 2000; Yang et al., 2013). Due to high translational activity in *E. coli*, cloning of plasmid-borne G10L-driven plastid transgenes is often complicated by recovery of sequence polymorphisms and structural rearrangements that attenuate the toxic levels of recombinant protein expression.

To test repression of transgenes whose expression is enhanced by the G10L, we designed two different gRNAs to target this element, as can be seen in Fig. 2C. The 27 nucleotide (nt) gRNAs were designed to target overlapping G10L sequences (gRNA7, gRNA8), and both targets include the YTTN protospacer adjacent motif and are surrounded by the constant repeat sequences required for gRNA recognition (Lin et al. 2021; Price et al., 2020). Each gRNA was driven by a constitutive P_tet_ promoter (Bertram and Hillen, 2008) and was integrated into strain PCP6 at the λ *attB* (Fig. 2B) site to create strains PCP7 and PCP8. The PCP7 and PCP8 strains carrying the P_rha_::dMad7 cassette and an individual gRNA were each confirmed via whole genome sequencing at both the φ80 and λ *attB* sites.

#### CRISPRi-mediated knockdown of transgene expression

To test the efficacy of the CRISPRi system, PCP7 and PCP8 strains were transformed with the high copy number plasmid, pPTS389, expressing GmP_rrn__G10L::GFP, to create PCP7_389 and PCP8_389 strains. The strains were then grown overnight in either the presence or absence of 1 mM rhamnose to measure attenuation of GFP fluorescence, anticipated after induction of gRNA expression.

As can be seen in Fig. 1, the PCP7_389 strain showed very high GFP fluorescence in the absence of rhamnose. In contrast, growth in the presence of rhamnose decreased GFP fluorescence intensity by at least six-fold (*p* < 0.0001). As control, the PCP2i strain carrying only an integrated dMad7 cassette and no gRNAs that was also transformed with pPTS389 (strain 2i_389, Fig. 1) and had high levels of GFP fluorescence in both the presence and absence of rhamnose. As expected, the PCP7 strain without the pPTS389 plasmid did not produce any GFP signal. These results indicate that dMad7 expression alone is not sufficient for the CRISPRi activity and that the inclusion of the guide RNA is required for knockdown of GFP fluorescence.

Growth of the PCP8_389 strain in overnight liquid culture was severely compromised independent of the presence or absence of rhamnose, apparently due to the toxic effect of GFP expression in this strain. As a result, GFP fluorescence intensity was measured only near background levels (Fig. 1). As a secondary approach to evaluate the CRISPRi system, agar medium plates were used to grow the bacterial strains, as shown in Fig. 3. In this case, PCP7 and PCP8 were transformed with plasmid pPTS522 carrying a GFP expression cassette driven by the soybean plastid *rbcL* gene promoter (GmP_rbcL_) and G10L (GmP_rbcL__G10L::GFP) to create PCP7_522 and PCP8_522 strains. While GmP_rbcL_ is also active in *E. coli*, its expression is weaker than GmP_rrn__G10L and often allows growth of recombinant strains. These new strains were then grown on agar plates with or without rhamnose and visualized under UV light after overnight growth. As in the result from liquid culture, the PCP7_522 had high GFP fluorescence in the absence of rhamnose and nearly abolished GFP signal in the presence of rhamnose. In contrast, the PCP8_522 strain showed significant GFP fluorescence in both the presence and absence of rhamnose in the agar medium. These results suggest that the gRNA8 in the PCP8_522 strain is much less effective and does not appreciably knockdown GFP expression, highlighting the importance of gRNA design in this transgene repression system and the success of the specific gRNA7.

**Figure 3.**
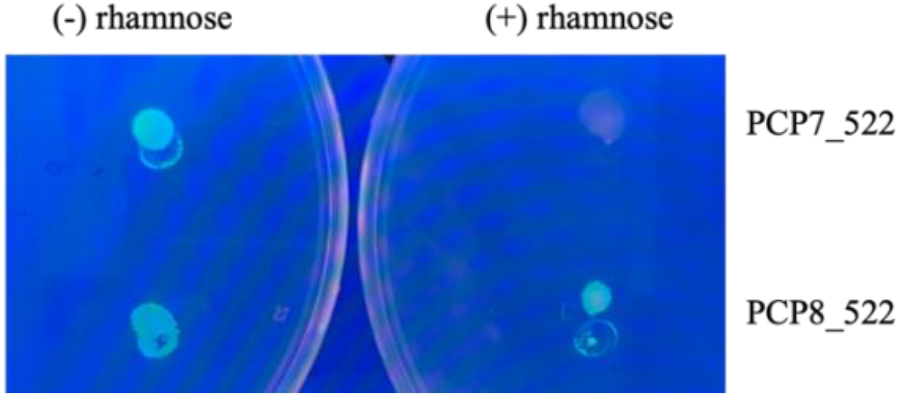
Visualization of strain GFP fluorescence on agar media plates. Strains PCP7 and PCP8 was transformed with pPTS522 and spotted on plates with or without rhamnose, grown overnight, and visualized under UV light. PCP7_522 shows a clear reduction in GFP signal in the presence of rhamnose, while PCP8_522 does not.

### HFQ approach to plasmid-borne toxic transgene repression Components of the Hfq transgene repression system

The CRISPRi approach described above was used successfully to repress plasmid-borne toxic transgene expression using an engineered dMad7 transgene integrated into the *E. coli* genome. However, this approach could be simplified further if an endogenous *E. coli* gene repression system were used as an alternative. To address this limitation, we have utilized a method of transgene repression based on the endogenous bacterial Hfq protein and a partner synthetic small RNA (Na et al., 2013; Sauer et al., 2012; Yoo et al., 2013) to knockdown the expression of our plasmid-borne toxic transgenes. While the Hfq system has been used to reduce the expression of some endogenous single copy *E. coli* genes (Yoo et al., 2013; Xie et al., 2020), this system has not been used to regulate expression of heterologous transgenes on multi-copy plasmids used in cloning applications.

The endogenous Hfq protein recognizes a synthetic sRNA when fused to an RNA scaffold encoded by the *E. coli* genome. Binding of Hfq protein to the sRNA-scaffold complex then triggers scanning to the target sequence in a transcript if the binding affinity is adequate. The Hfq sRNA-scaffold complex then impedes translation of the mRNA and causes degradation of the transcript through the recruitment of RNaseE (Dwijayanti et al., 2022), reducing expression of the encoded protein. In our repression system, we have engineered the sRNA to recognize a plastid transgene sequence of interest, enabling the Hfq-sRNA scaffold complex to target heterologous transgenic mRNA sequences. Since Hfq and the sRNA scaffold are endogenous bacterial elements, only the sRNA needs to be engineered to recognize the target of interest.

#### Design of engineered sRNA targets

We designed sRNA sequences of between 20-30 nucleotides with a Gibbs Free Energy of binding (Δ*G*_*bind*_) calculated by DuplexFold software (Reuter and Mathews, 2010) of -35 kcal/mol considered optimal for target recognition (Na et al., 2013). The engineered sRNAs were fused to the 79 nt scaffold from the *E. coli micC* coding sequence (Yoo et al., 2013). We targeted the G10L translational enhancer, to compare the efficacy of the Hfq repression approach to the previous CRISPRi results. Additional sRNA targets were designed against the *aadA* selectable marker and GFP coding regions also typically used in plastid transgenic plants to test the versatility and modularity of this system (Fig. 4).

**Figure 4.**
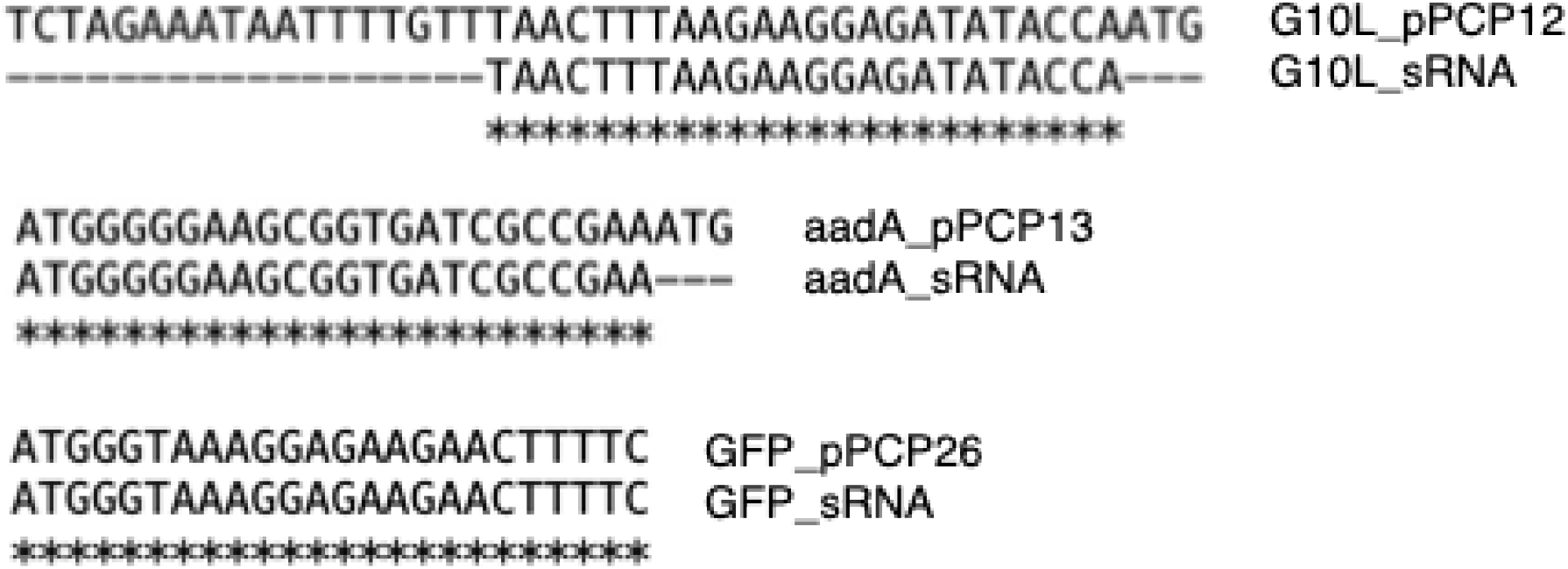
sRNA sequences and alignment to their targets. Target sequences are labeled with an underscore and the plasmid name that harbors that control sequence whereas the small RNA sequences are labeled with an underscore and sRNA. Asterisks indicate sequence identities.

To further expand the utility of this repression approach, a triplex sRNA-scaffold was also created to target all three transgene elements simultaneously (Supplemental Fig. 1). In this case, the three specific sRNA-scaffolds were arranged in tandem, each flanked by either HDV or hammerhead ribozymes designed to self-splice and release the individual sRNA-scaffolds from the triplex sRNA transgene, an approach that has been successful for processing of CRISPR gRNAs in vitro and in yeast (Gao & Zhao, 2014; Lin et al. 2021) but has not been tested for larger sRNA-scaffold sequences used in the Hfq system.

#### Efficacy testing of the triplex sRNA-scaffolds

To test the efficacy of the triplex sRNA-scaffold sequences, these were expressed from both high copy number plasmid-based testing constructs and via a triplex cassette integrated into the *E. coli* genome. Plasmids pPCP12 and pPCP13 (Supplemental Fig. 1) each carry the triplex sRNA-scaffold cassette expressed from the strong bacterial promoter, J23119 (https://parts.igem.org/Part:J23119; Anderson, J.C., promoter collection) and a nanoluciferase reporter transgene with an upstream target sequence of interest. In pPCP12, the nanoluciferase coding region is expressed from the *E. coli* J23105 promoter (https://parts.igem.org/Part:J23105; Anderson, J.C., promoter collection;) and has as target a 42 nt G10L translation enhancer sequence downstream of the transcription start site. In pPCP13, the promoter of soybean plastid *psbA* gene (GmP_psbA_), that has strong transcriptional activity in *E. coli* (Cohen et al., 1984; Efimov et al., 1994), drives expression of a nanoluciferase gene carrying as repression target an N-terminal fusion of the first 8 amino acids of the *aadA* coding region.

*E. coli* NEB Stable cells were transformed with the pPCP12 and pPCP13 plasmids, and stable recombinant strains (PCP12 and PCP13, respectively) were confirmed by plasmid sequencing. The PCP12 and PCP13 strains were then grown along with a strain carrying the positive control plasmid, pSCP6040, that carries the identical nanoluciferase reporter gene as in pPCP12. Reduction of luciferase activity in the overnight cultures relative to control is indicative of successful ribozyme cleavage of the triplex sRNA-scaffolds and release of individual efficacious sRNA-scaffolds that bind to their cognate target to inhibit luciferase activity. As can be seen in Fig. 5A, luciferase activity was significantly decreased by ∼65% when either the G10L or *aadA* sequence was targeted by the sRNA-scaffolds, indicating successful repression of the transgene sequences in this approach. Furthermore, since the reporter transgenes are carried on high copy number plasmids and expressed from a strong promoter, the level of target knockdown appears to be surprisingly very efficient.

**Figure 5.**
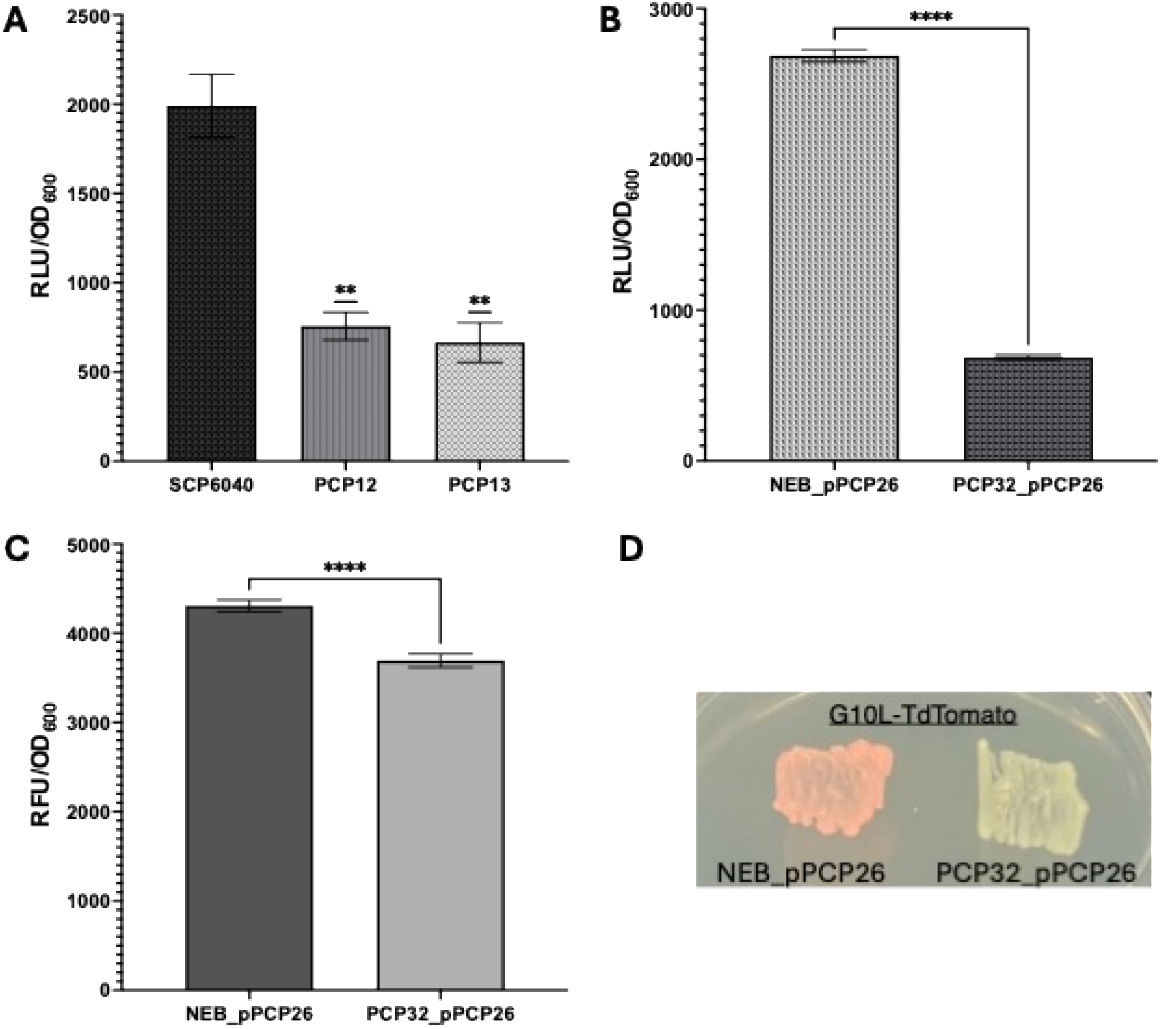
Triplex sRNA-scaffold knockdown of target mRNAs. Cultures were grown overnight and then diluted with LB back to OD600 = 0.001 prior to the assay. **A)** Strains PCP12 and PCP13 carry a luciferase reporter cassette containing either G10L or aadA sRNA target sequences, respectively, and the corresponding sRNA-scaffold on a high-copy plasmid. Strain SCP6040 carries the same luciferase cassette without sRNA target or scaffold sequences. **B)** A triplex sRNA-scaffold was integrated into the *E. coli* genome to generate strain PCP32, which was then transformed with plasmid pPCP26 carrying three reporter genes to evaluate repression. Repression of the aadA reporter reached ∼60%. **C)** Repression of the GFP reporter by the GFP sRNA-scaffold and target sequence was ∼15%. **D)** Complete repression of TdTomato by the G10L sRNA-scaffold and target sequence is evident from the absence of pink pigment in bacteria grown on agar. Plasmid pPCP26 transformed into the NEB strain lacking an integrated sRNA-scaffold served as the control.

#### Efficacy testing of an integrated copy of the triplex sRNA-scaffold

Integration of the sRNA multiplex expression cassette was achieved using CRIM system (Haldimann and Wanner, 2001) as described above. NEB Stable cells were transformed with the pMAX851 *ϕ*80 integrase helper plasmid and then subsequently with a plasmid that contains the triplex sRNA-scaffold expression cassette designed for integration at *E. coli ϕ*80 *attB* site (Supplemental Fig. 2). The helper plasmid was then cured by overnight growth at 42 ºC to form strain PCP32. The PCP32 strain was verified to be base-perfect by whole genome sequencing.

The PCP32 strain carrying the integrated triplex sRNA-scaffold expression cassette was then transformed with a multi-reporter high copy number plasmid, pPCP26, for simultaneous efficacy testing of sRNA-scaffolds. Plasmid pPCP26 carries three reporter genes (TdTomato, GFP and nanoluciferase), that are fused to a target sequence of interest (Supplemental Fig. 1), either in the 5’-untranslated G10L (ZmPrrn_G10L::TdTomato), as translational fusion (aadA-nanoluciferase), or the native GFP N-terminal sequence, such that Hfq-mediated repression via the target sequence should inhibit reporter gene expression.

As can be seen in Fig. 5B, the expression of the nanoluciferase reporter gene that is translationally fused to an *aadA* target sequence was dramatically reduced by more than 60% in this assay. As shown in Fig. 5C, the GFP reporter expression was also significantly reduced when its first 8 amino acids is used as the Hfq target, although repression in that case was only a moderate ∼15%, despite both sRNAs being 100% complementary to their target sequences (Fig. 4). It’s possible that exceptional stability of the GFP protein (Andersen et al., 1998) contributes to maintenance of activity in this fluorescence assay, while visible fluorescence was dramatically reduced when observed from bacterial cultures on agar medium in the CRISPRi approach (Fig. 3).

A quantitative assay was not available to measure TdTomato repression mediated by the G10L sequence in plasmid pPCP26. However, high TdTomato expression in unmodified *E. coli* NEB Stable cells (NEB_pPCP26) results in an intense pink color in cultures, as shown in Fig. 5D. In contrast, as previously observed in the CRISPRi approach, TdTomato repression via the G10L target was very efficient, completely eliminating visible pink G10L::TdTomato expression in the PCP32 strain carrying the pPCP26 plasmid (PCP32_pPCP26).

#### Improved cloning outcomes using the Hfq transgene repression system

The efficacy of the Hfq system was further tested via use of the PCP32 strain as host for standard cloning vector assembly compared to the commercial NEB Stable cells used as bacterial host. As a test case, the results from a four-piece Gibson assembly of the pPCP26 multi-reporter plasmid described above were used to examine the recovery of sequence polymorphisms or structural rearrangements when the PCP32 strain was used as the cloning host, compared with unmodified NEB Stable cells as host. After transformation of each host with the identical 4-fragment Gibson reaction, recombinant colonies were randomly chosen for sequence analysis to characterize any sequence polymorphisms or structural rearrangements.

As shown in Fig. 6, deletion mutations and sequence polymorphism were recovered in the colonies derived from the unmodified NEB Stable cells host. In contrast, no mutations were recovered, and the 4-fragment cloning was correctly assembled, in all colonies derived from the Hfq-engineered PCP32 strain as host. Several colonies derived from both strains varied by a single nucleotide in a poly A sequence tract that is likely due to sequencing artifacts rather than actual cloning errors. These results confirm the efficacy of the engineered Hfq system to minimize random mutations caused by toxic overexpression of recombinant proteins.

**Figure 6.**
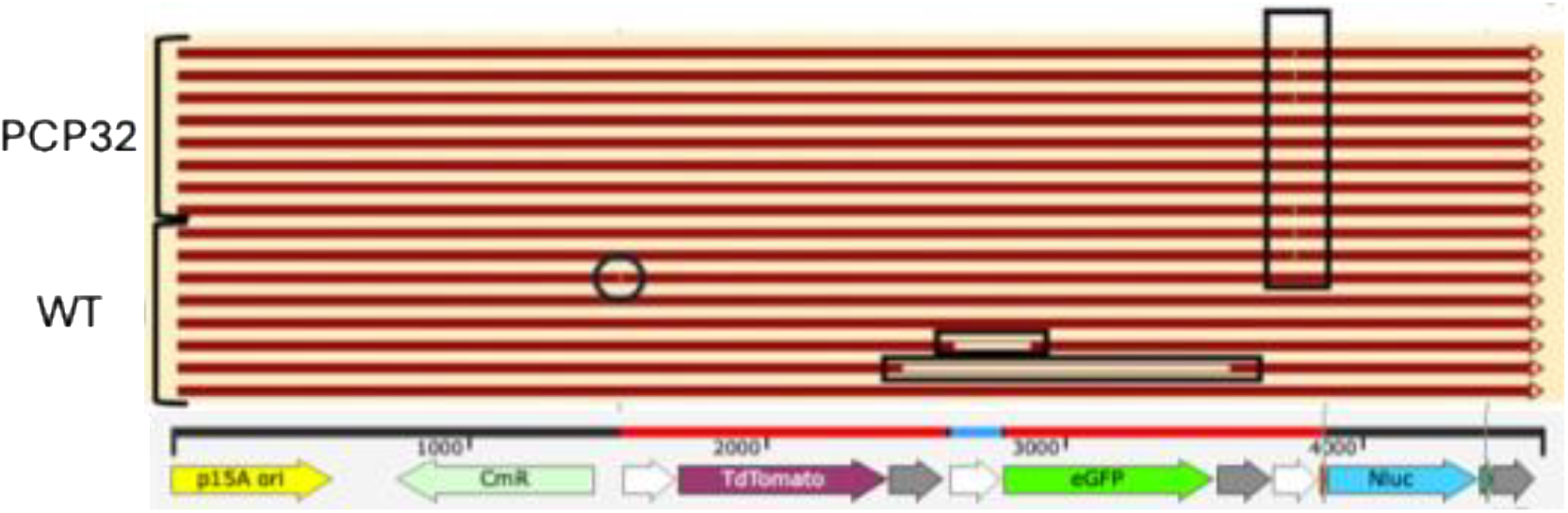
Sequencing results of independently transformed colonies derived from the 4-way assembly of plasmid pPCP26 in the NEB Stable cell background (WT, bottom bracket) versus the Hfq-engineered PCP32 strain (top bracket) that has the sRNA multiplex expression cassette integrated into the genome. Note the lack of any mutations in eight sequenced clones from the PCP32 strain versus three of the eight clones sequenced from the NEB Stable cells that contained mutations preventing use of the plasmid. Horizontal black boxes and circle show the locations of deletions and a single nucleotide mutation in independent colonies from transformation into the NEB strain. The vertical black box shows the position of the polyA tract that has some sequence ambiguities. CmR, chloramphenicol resistance; Nluc, nanoluciferase.

## DISCUSSION

As the field of synthetic biology grows increasingly complex, so must the tools improve to support expression of larger and more ambitious transgene circuits. Cloning has long been a bottleneck due to detrimental effects on *E. coli* physiology, the organism most likely to be the production platform, that result in rearrangement and mutation of transgene sequences to attenuate their expression. These complexities are exacerbated in cases where transgene expression in the cloning host is unavoidable, as in engineering of prokaryotic-like organelles, chloroplasts and mitochondria. Described here are two different transgene repression systems that enabled cloning of unavoidably highly expressed and potentially toxic plastid transgenes on high copy number plasmids in *E. coli*, where a commercial bacterial strain was unable to do the same. Knockdown of plastid transgenes was demonstrated using observable reporter readouts, such as luciferase activity and fluorescent protein accumulation. Substantial knockdowns were measured from both systems: as much as a six-fold reduction in gene expression from the CRISPRi system and between 15 - 65% reduction from the Hfq system. Our CRISPRi and Hfq approaches extend previous work with individual toxic genes (Wen, et al., 2021) and endogenous single copy *E. coli* genes (Yoo et al., 2013; Xie et al., 2020), to a multiplex system that can simultaneously repress expression of at least three highly expressed transgenes on multicopy plasmids.

The CRISPRi system required inducible expression and an integrated single copy of dMad7 to keep its uninduced expression low to avoid potential toxicity to the host cell (Mund et al., 2023). Interestingly, we could only recover bacterial strains that cointegrated a native insertion element next to dMad7. Although rhamnose inducible knockdown of multiple plasmid-bone transgenes was still efficient from this system (Fig. 1), it’s possible that the insertion element further limited dMad7 expression in ways favorable to bacterial growth. Moving forward, dMad7 expression could be optimized to avoid toxicity but enhance efficacy via testing of additional inducible or titratable promoters. On the other hand, the endogenous Hfq protein only requires a single engineered small RNA component for functionality, unlike other approaches that require multiple pieces (Vandierendonck et al., 2023). Further, the Hfq system is inducer free, programmable for any sequence target and has no toxic components like the CRISPR-based system. Like the CRISPRi system, Hfq was also efficient at repression of transgenes carried on high copy number plasmids. Sequence and structural integrity of a 4-piece ligation plasmid was recovered in multiple independent plasmid clones (eight of eight), whereas various mutations accumulated in several (three of eight) plasmid clones in the absence of the sRNA multiplex expression cassette. Despite the small sample size, this result shows promise in employing the customized Hfq system for problematic clonings. Recent work in our lab (data not shown) confirms that even DNA breakage enzymes that are known to be toxic to *E. coli* when overexpressed are now efficiently cloned using Hfq-based transgene silencing.

The multiplex sRNA-scaffolds were apparently efficiently processed into individual functional units by flanking self-processing ribozymes, as evidenced by their efficacy in transgene repression. Thus, the upper limit of engineered scaffolds and target sequences is yet to be determined. In both the CRISPRi and Hfq systems, there was no apparent loss of plasmid yields when three expressible transgenes were present. This also suggests that the artificial sRNAs had no significant genomic off-targets, indicating that increased expression of the sRNA-scaffolds may also be another approach to enhanced efficacy of the approaches. Future system designs could match sRNA expression to the perceived level of toxicity of the target genes. To this end, our modular triplex sRNA-scaffold cassette separates its promoter and individual gene sRNA-scaffolds by unique restriction enzyme sites (Supplemental Fig. 1) for easy sequence exchange directly into the backbone of a high copy number plasmid.

Here we have presented two transgene repression systems to address cloning of toxic and unstable constructs. Data for the repression of transgenes in the cloning host indicate clear knockdown of transgenic sequences, as well as the improvement of cloning outcomes due to the engineered systems. Our primary application of these systems is to clone unavoidably expressed transgenes in constructs destined for plant chloroplast engineering; however, the scope of possible applications is broad as evidenced by the repression of GFP and *aadA*, which are common transgenic elements found in many areas of bioengineering. Therefore, these applications of CRISPRi and Hfq are poised to make an impact in the field of synthetic biology by streamlining the creation of difficult constructs, enabling advances that might not otherwise be achievable.

## MATERIALS AND METHODS

### Strain growth

A complete list of plasmids and bacterial strains made during this work is listed in Supplemental Figure 3. Bacterial strains were grown in LB media; antibiotic additions were adjusted based on the copy number of the drug resistance cassette. For plasmid growth, the following concentrations were used (all values are μg/mL): ampicillin 125, tetracycline 12.5, kanamycin 50 and chloramphenicol 25; for chromosomal inserts: tetracycline 8 and kanamycin 25. For the induction of the P_rha_ reporter, rhamnose was added to a final concentration of 1 mM. All cultures were grown at 37 ◦C.

### Strain construction and verification

All plasmids were constructed using a Gibson assembly approach (Gibson et al., 2009). All strains were confirmed via Oxford Nanopore Sequencing. Bacterial genome integration strains were verified by amplicon sequencing, and by whole genome sequencing (Plasmidsaurus). Integration into the *E. coli* chromosome of dMad7 CDS, individual guide RNAs and Hfq triplex seed RNA cassettes was achieved using the conditional-replication, integration, and modular (CRIM) system developed by Haldimann and Wanner (2001).

Integration of dMad7 and gRNA expression cassettes:

Plasmid pPCP2 carries the dMad7 expression cassette driven by a rhamnose inducible promoter, P_rha_. pPCP2 has an R6K replication origin that is dependent upon the pi protein for replication and is maintained in *pir+* strains for propagation (Metcalf et al., 1994). DH10B cells (*pir-*) were transformed with plasmid pMAX851, a φ80 integrase helper plasmid with temperature sensitive origin of replication, to create the DH10B_tsSC101 strain. DH10B_tsSC101 cells were made electrocompetent and transformed with the pPCP2 plasmid carrying the P_rha_::dMad7 expression construct. Growth at 37 ◦C allowed for integration of P_rha_::dMad7 at the φ80 *attB* site, after which the helper plasmid was subsequently cured from the recombinant strain by overnight growth at 42 ◦C, to form the PCP2i strain. PCP2i cells were then made electrocompetent and transformed with a λ integrase helper plasmid, pMAX853, to form PCP6. PCP6 was then made competent and transformed with each of the gRNA plasmids, pPCP3 and pPCP4; integration of the gRNA and subsequent curing of the λ integrase helper plasmid was conducted as before. The resulting strains – PCP7 and PCP8 – were used in downstream experiments.

Integration of the triplex sRNA-scaffold. expression cassette:

NEB Stable cells were transformed with plasmid pMAX851 to create strain NEB_tsSC101. NEB_tsSC101 cells were made electrocompetent and transformed with pPCP24, which contains the triplex seed RNA expression cassette in a vector designed for integration at the bacterial *ϕ*80 *attB* integration site. Initial growth at 37 ºC allowed for integration at the *ϕ*80 *attB* site; the helper plasmid was then subsequently cured by overnight growth at 42 ºC to form PCP32 strain. PCP32 cells were then made competent and transformed with plasmid pPCP26 to form strain PCP32_26 carrying a triplex expression reporter for the knockdown experiments.

### GFP and Luciferase assays

Cells were grown overnight at 37 ◦C to an OD_600_ > 2 with appropriate antibiotics and in either the presence or absence of rhamnose. These saturated cultures were diluted back to OD_600_ = 0.001, then aliquoted (100 μL) in triplicate into the wells of a clear bottom 96 well plate (Costar). Luciferase activity was measured after the addition of substrate (NanoDLR Stop & Glo, Promega) in a 96 well plate reader, fitted with photomultiplier tubes (POLARstar Omega, BMG Laboratories). GFP activity was measured as light emission at 509λ, following excitation at 395λ. This value was normalized to the culture density for a final value of RFU/OD_600_. To visualize GFP knockdown on agarose plates, UV light was used to excite GFP, which was visualized in a gel imager (Azure Biosystems).

## Author Contributions

JMS and AP conceived of the CRISPRi studies, AP conceived of the Hfq studies, AP conducted all the experimental work, and JMS and AP wrote the manuscript.

## Funding and Acknowledgments

All work was funded by Plastomics Inc. We thank all members of Plastomics laboratory for their support during this work, including Kara Boltz, Scott Dour, Alice Hui and Zuzana Kocsisova for helpful discussions and advice.

## Availability of Materials

Plasmids and strains detailed in this publication may be available under Material Transfer Agreement to non-profit academic researchers.

