## Supplemental files for "Small RNA-guided transgene repression systems enable toxic gene cloning in bacteria"

**SUPPLEMENTAL FIGURES**

**Supplemental Figure 1**

>pPCP24 triplex sRNA-scaffolds

**5’TTGACAGCTAGCTCAGTCCTAGGTATAATGCTAGCACCGGT**ATACCACTGATGAGTCCGTGAGGACGAAACGAGTAAGCTCGTCTGGTATATCTCCTTCTTAAAGTTATTTCTGTTGGGCCATTGCATTGCCACTGATTTTCCAACATATAAAAAGACAAGCCCGAACAGTCGTCCGGGCTTTTTTT**GCCGGCATGGTCCCAGCCTCCTCGCTGGCGCCGGCTGGGCAACATGCTTCGGCATGGCGAATGGGAC**cgccggcgtagaaaggggccggccCTTTTCCTGATGAGTCCGTGAGGACGAAACGAGTAAGCTCGTC**GAAAAGTTCTTCTCCTTTACCCAT**TTTCTGTTGGGCCATTGCATTGCCACTGATTTTCCAACATATAAAAAGACAAGCCCGAACAGTCGTCCGGGCTTTTTTT**GGCCGGCATGGTCCCAGCCTCCTCGCTGGCGCCGGCTGGGCAACATGCTTCGGCATGGCGAATGGGAC**tccggatacgaaggcaacacgcgtGCCGAACTGATGAGTCCGTGAGGACGAAACGAGTAAGCTCGTC**TTCGGCGATCACCGCTTCCCCCAT**TTTCTGTTGGGCCATTGCATTGCCACTGATTTTCCAACATATAAAAAGACAAGCCCGAACAGTCGTCCGGGCTTTTTTT**GGCCGGCATGGTCCCAGCCTCCTCGCTGGCGCCGGCTGGGCAACATGCTTCGGCATGGCGAATGGGACCGTACGGTCAGTTTCACCTGTTTTACGTAAAAACCCGCTTCGGCGGGTTTTTACTTTTGG 3’**

J23119 promoter, bold black; HH ribozyme, red; G10L sRNA, blue; micC scaffold, green; HDV ribozyme, bold green; spacer sequences, lowercase; GFP sRNA, bold purple; aadA sRNA sequence, bold orange; garPLRK-rnpB terminator, bold black. Restriction enzyme sites are underlined.

>pPCP12 triplex sRNA-scaffold with G10L target sequence upstream of nanoluciferase

**5’TTGACAGCTAGCTCAGTCCTAGGTATAATGCTAGCACCGGT**ATACCACTGATGAGTCCGTGAGGACGAAACGAGTAAGCTCGTCTGGTATATCTCCTTCTTAAAGTTATTTCTGTTGGGCCATTGCATTGCCACTGATTTTCCAACATATAAAAAGACAAGCCCGAACAGTCGTCCGGGCTTTTTTT**GCCGGCATGGTCCCAGCCTCCTCGCTGGCGCCGGCTGGGCAACATGCTTCGGCATGGCGAATGGGAC**cgccggcgtagaaaggggccggccCTTTTCCTGATGAGTCCGTGAGGACGAAACGAGTAAGCTCGTC**GAAAAGTTCTTCTCCTTTACCCAT**TTTCTGTTGGGCCATTGCATTGCCACTGATTTTCCAACATATAAAAAGACAAGCCCGAACAGTCGTCCGGGCTTTTTTT**GGCCGGCATGGTCCCAGCCTCCTCGCTGGCGCCGGCTGGGCAACATGCTTCGGCATGGCGAATGGGAC**tccggatacgaaggcaacacgcgtGCCGAACTGATGAGTCCGTGAGGACGAAACGAGTAAGCTCGTC**TTCGGCGATCACCGCTTCCCCCAT**TTTCTGTTGGGCCATTGCATTGCCACTGATTTTCCAACATATAAAAAGACAAGCCCGAACAGTCGTCCGGGCTTTTTTT**GGCCGGCATGGTCCCAGCCTCCTCGCTGGCGCCGGCTGGGCAACATGCTTCGGCATGGCGAATGGGACCGTACGGTCAGTTTCACCTGTTTTACGTAAAAACCCGCTTCGGCGGGTTTTTACTTTTGG**ggatccccgggtaccgagctcgaattctcagtcgttgtcgttagcagaaagtcaaaagcctccgaccggaggcttttgact**TTTACGGCTAGCTCAGTCCTAGGTACTATGCTAGCTATAGTCTAGAAATAATTTTGTTTAACTTTAAGAAGGAGATATACCA**ATGGTATTTACTTTAGAAGATTTTGTAGGAGATTGGAGACAAACTGCTGGATATAATTTAGATCAAGTATTAGAACAAGGAGGAGTATCTTCTTTATTTCAAAATTTAGGAGTATCTGTAACTCCTATTCAAAGAATTGTATTATCTGGAGAAAATGGATTGAAAATTGATATTCATGTAATCATTCCTTATGAAGGATTATCTGGAGATCAAATGGGACAAATTGAAAAGATCTTTAAAGTAGTATATCCTGTTGATGATCATCATTTCAAAGTAATCTTGCATTATGGAACTTTAGTAATAGATGGTGTTACACCAAACATGATTGATTATTTCGGTCGTCCATATGAGGGTATAGCAGTTTTCGATGGTAAGAAAATAACTGTTACTGGAACACTTTGGAATGGTAATAAGATTATAGATGAACGTCTTATAAATCCTGATGGAAGTCTTTTGTTTAGAGTAACTATAAATGGAGTTACAGGATGGAGATTATGTGAAAGAATTTTAGCTTAA

J23119 promoter, bold black; HH ribozyme, red; G10L sRNA, blue; micC scaffold, green; HDV ribozyme, bold green; spacer sequences, lowercase; GFP sRNA, bold purple; aadA sRNA sequence, bold orange; garPLRK-rnpB terminator, bold black; spacer sequences, lowercase; J23105 promoter, bold black; G10L target sequence, bold blue; Nanoluciferase, magenta. Restriction enzyme sites are underlined.

>pPCP13 triplex sRNA-scaffold with aadA target sequence upstream of nanoluciferase

**5’TTGACAGCTAGCTCAGTCCTAGGTATAATGCTAGCACCGGT**ATACCACTGATGAGTCCGTGAGGACGAAACGAGTAAGCTCGTCTGGTATATCTCCTTCTTAAAGTTATTTCTGTTGGGCCATTGCATTGCCACTGATTTTCCAACATATAAAAAGACAAGCCCGAACAGTCGTCCGGGCTTTTTTT**GCCGGCATGGTCCCAGCCTCCTCGCTGGCGCCGGCTGGGCAACATGCTTCGGCATGGCGAATGGGAC**cgccggcgtagaaaggggccggccCTTTTCCTGATGAGTCCGTGAGGACGAAACGAGTAAGCTCGTC**GAAAAGTTCTTCTCCTTTACCCAT**TTTCTGTTGGGCCATTGCATTGCCACTGATTTTCCAACATATAAAAAGACAAGCCCGAACAGTCGTCCGGGCTTTTTTT**GGCCGGCATGGTCCCAGCCTCCTCGCTGGCGCCGGCTGGGCAACATGCTTCGGCATGGCGAATGGGAC**tccggatacgaaggcaacacgcgtGCCGAACTGATGAGTCCGTGAGGACGAAACGAGTAAGCTCGTC**TTCGGCGATCACCGCTTCCCCCAT**TTTCTGTTGGGCCATTGCATTGCCACTGATTTTCCAACATATAAAAAGACAAGCCCGAACAGTCGTCCGGGCTTTTTTT**GGCCGGCATGGTCCCAGCCTCCTCGCTGGCGCCGGCTGGGCAACATGCTTCGGCATGGCGAATGGGACCGTACGGTCAGTTTCACCTGTTTTACGTAAAAACCCGCTTCGGCGGGTTTTTACTTTTGG**ggatccccgggtaccgagctcgaattctcagtcgttgtcgttagcagaaagtcaaaagcctccgaccggaggcttttgact**GAGGTAAAAAAGTAAATTCGGATAAATCTAAATAAGAGCATACTTACTATGGATATTGGTATTGGTTGACACTGGTATATAAGTCATGTTATACTGTTGAATAACAAGTCCTCAAATTTTCTAATTCTAGATAATTTTGGTGCTTGGGAGTCCCTAAAGATTAAATCAACCAAGATTTTACAATGGGGGAAGCGGTGATCGCCGAA**ATGGTATTTACTTTAGAAGATTTTGTAGGAGATTGGAGACAAACTGCTGGATATAATTTAGATCAAGTATTAGAACAAGGAGGAGTATCTTCTTTATTTCAAAATTTAGGAGTATCTGTAACTCCTATTCAAAGAATTGTATTATCTGGAGAAAATGGATTGAAAATTGATATTCATGTAATCATTCCTTATGAAGGATTATCTGGAGATCAAATGGGACAAATTGAAAAGATCTTTAAAGTAGTATATCCTGTTGATGATCATCATTTCAAAGTAATCTTGCATTATGGAACTTTAGTAATAGATGGTGTTACACCAAACATGATTGATTATTTCGGTCGTCCATATGAGGGTATAGCAGTTTTCGATGGTAAGAAAATAACTGTTACTGGAACACTTTGGAATGGTAATAAGATTATAGATGAACGTCTTATAAATCCTGATGGAAGTCTTTTGTTTAGAGTAACTATAAATGGAGTTACAGGATGGAGATTATGTGAAAGAATTTTAGCTTAA

J23119 promoter, bold black; HH ribozyme, red; G10L sRNA, blue; micC scaffold, green; HDV ribozyme, bold green; spacer sequences, lowercase; GFP sRNA, bold purple; aadA sRNA sequence, bold orange; garPLRK-rnpB terminator, bold black; spacer sequences, lowercase; J23105 promoter, bold black; GmPpsbA promoter, bold red; aadA target sequence, bold light blue; Nanoluciferase, magenta. Restriction enzyme sites are underlined.

>pPCP26 with three reporter genes controlled by upstream sRNA targets

**CCACGATCGAACGGGAATGGATAGGAGGCTTGTGGGATTGACGTGATAGGGTAGGGTTGGCTATACTGCTGGTGGCGAACTCCAGGCTAATAATCTGAAGCGCATGGATACAAGTTATCCTTGGAAGGAAAGACAATTCCGGATCCTGTAGAAATAATTTTGTTTAACTTTAAGAAGGAGATATA**CCCATGGTTTCTAAAGGAGAAGAAGTTATTAAAGAATTTATGAGATTTAAAGTTAGAATGGAAGGATCTATGAATGGACATGAATTTGAAATTGAAGGAGAAGGAGAAGGTAGACCTTATGAAGGAACTCAAACTGCTAAATTAAAAGTTACTAAAGGAGGACCTTTACCTTTTGCTTGGGACATTTTAAGTCCTCAATTTATGTATGGATCTAAAGCTTATGTTAAACATCCTGCTGACATTCCTGACTATAAGAAGTTATCTTTTCCTGAAGGATTTAAATGGGAAAGAGTTATGAATTTTGAAGACGGAGGTTTAGTTACTGTAACTCAAGACTCCAGTTTACAAGACGGAACACTTATTTATAAAGTAAAGATGAGAGGAACTAATTTTCCTCCAGATGGTCCAGTTATGCAAAAGAAGACAATGGGATGGGAAGCTTCTACTGAGCGTCTTTATCCTCGTGACGGTGTATTAAAAGGTGAGATTCATCAAGCTCTTAAACTTAAAGATGGAGGACACTATTTAGTAGAATTCAAAACTATATATATGGCAAAGAAACCAGTACAATTACCTGGATATTATTATGTTGACACTAAACTTGACATAACATCTCATAATGAGGACTATACTATTGTCGAACAATATGAGAGATCTGAAGGAAGACATCATCTTTTCTTATATGGTATGGACGAGCTTTATAAGTAG**TTTTTAAATTGATTCAATTGTGAAATAACACGACATGTGTATCTAGGGAATAGTTTCTTCAAAGCGAATTCTCCCTAGATACATCTATTCAATTTAATTCTGAATTTATTTTGAATATATGATATATTAATATATTAATTGTGCTAAAGAGTTTCAATCTATTTTCACTAAGTAAGTCCAATAG**ctcgagggggggcgggg**CCCAAATTTTTGGATTTGGTAAATGAAGTTATACGAAAATCCAATCGTTGGGGCTGGCTTGGTTGACATTGGTATATAGACTATGTTATACTGTTAAATAACAAGCCTTCTATTATCTATTTTCTTTCTAGTTAATACGTGTGCTTGGGAGTCCTTGCAATTTGAATAAACCAAGATCTTACC**ATGGGTAAAGGAGAAGAACTTTTCACTGGAGTTGTCCCAATTCTTGTTGAATTAGATGGTGATGTTAATGGGCACAAATTTTCTGTCAGTGGAGAGGGTGAAGGTGATGCAACATACGGAAAACTTACCCTTAAATTTATTTGCACTACTGGAAAACTACCTGTTCCATGGCCAACACTTGTCACTACTTTCTGTTATGGTGTTCAATGCTTTTCAAGATACCCAGATCATATGAAGCGGCACGACTTCTTCAAGAGCGCCATGCCTGAGGGATACGTGCAGGAGAGGACCATCTTCTTCAAGGACGACGGGAACTACAAGACACGTGCTGAAGTCAAGTTTGAGGGAGACACCCTCGTCAACAGGATCGAGCTTAAGGGAATCGATTTCAAGGAGGACGGAAACATCCTCGGCCACAAGTTGGAATACAACTACAACTCCCACAACGTATACATCATGGCCGACAAGCAAAAGAACGGCATCAAAGCCAACTTCAAGACCCGCCACAACATCGAAGACGGCGGCGTGCAACTCGCTGATCATTATCAACAAAATACTCCAATTGGCGATGGCCCTGTCCTTTTACCAGACAACCATTACCTGTCCACACAATCTGCCCTTTCGAAAGATCCCAACGAAAAGAGAGACCACATGGTCCTTCTTGAGTTCGTAACAGCTGCTGGGATTACACATGGCATGGATGAACTAATCTAG**TTCGATTTTTAAATTGATTCAATTGTGAAATAACACGACATGTGTATCTAGGGAATAGTTTCTTCAAAGCGAATTCTCCCTAGATACATCTATTCAATTTAATTCTGAATTTATTTTGAATATATGATATATTAATATATTAATTGTGCTAAAGAGTTTCAATCTATTTTCACTAAGTAAGTCCAATAG**ACCCATTCGTAACAACTTTACTTTATTTAGTATTCCTTTTTTTATATTTAGTATTCCTAAAAAAAAAAATGCAATATAATAAAAATAAATCATTTTTAACACGATAAGCTAATTCTTACGTTTCCACACTAAAGTTAGATATAGTATTTTATTATTT**ATGGGGGAAGCGGTGATCGCCGAA**ATGGTATTTACTTTAGAAGATTTTGTAGGAGATTGGAGACAAACTGCTGGATATAATTTAGATCAAGTATTAGAACAAGGAGGAGTATCTTCTTTATTTCAAAATTTAGGAGTATCTGTAACTCCTATTCAAAGAATTGTATTATCTGGAGAAAATGGATTGAAAATTGATATTCATGTAATCATTCCTTATGAAGGATTATCTGGAGATCAAATGGGACAAATTGAAAAGATCTTTAAAGTAGTATATCCTGTTGATGATCATCATTTCAAAGTAATCTTGCATTATGGAACTTTAGTAATAGATGGTGTTACACCAAACATGATTGATTATTTCGGTCGTCCATATGAGGGTATAGCAGTTTTCGATGGTAAGAAAATAACTGTTACTGGAACACTTTGGAATGGTAATAAGATTATAGATGAACGTCTTATAAATCCTGATGGAAGTCTTTTGTTTAGAGTAACTATAAATGGAGTTACAGGATGGAGATTATGTGAAAGAATTTTAGCTTAA

ZmPrrn promoter, bold black; G10L target sequence, bold green; TdTomato, red; NtTpetD terminator, bold black; spacer, lowercase; ZmPpsbA promoter target sequence, bold blue; GFP, green; soybean plastid promoter, orange; aadA target sequence, bold orange; Nanoluciferase, magenta

**Supplemental Figure 2**

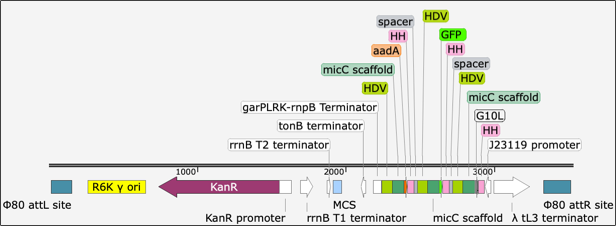

Triplex sRNA-scaffold integrated into the E. coli $\phi$80 site. The triplex is driven by the J23119 promoter and carries sRNA-scaffolds targeting aadA, GFP and G10L sequences. Each sRNA-scaffold is flanked by hammerhead (HH) and HDV ribozymes.

**Supplemental Figure 3**

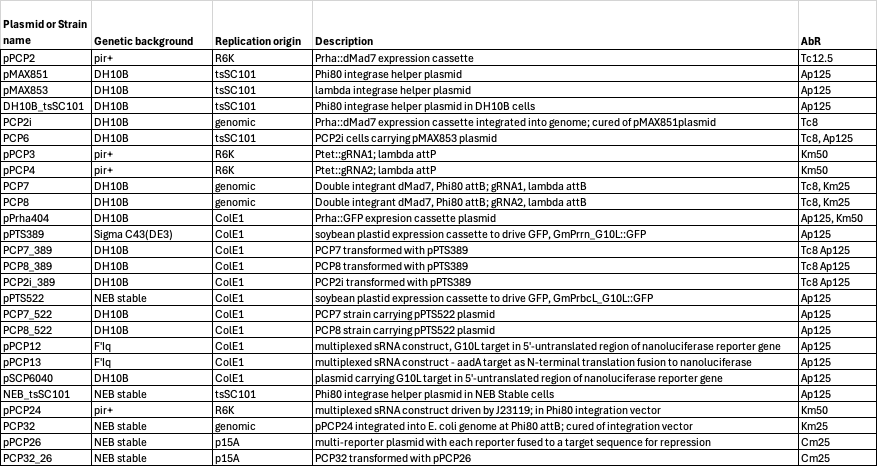

List of bacterial strains and plasmids used in this work
